# Citizen scientists highlight conservation value of a small subtropical reef, Flinders Reef, southeast Queensland, Australia

**DOI:** 10.1101/602631

**Authors:** Monique G.G. Grol, Julie Vercelloni, Tania M. Kenyon, Elisa Bayraktarov, Cedric P. van den Berg, Daniel Haris, Jennifer A. Loder, Morana Mihaljević, Phebe I. Rowland, Chris M. Roelfsema

**Author notes:** Corresponding author (MGGG). Author contributions These authors contributed equally to this work (Monique G.G. Grol, Julie Vercelloni, Tania M. Kenyon, Chris M. Roelfsema). Author contributions These authors also contributed equally to this work (Elisa Bayraktarov, Cedric P. van den Berg, Jennifer A. Loder, Morana Mihaljević, Phebe I. Rowland).

## Abstract

Subtropical reefs are unique ecosystems that require effective management – informed by regular ecological monitoring – to foster resilience to environmental changes. Resources to conduct monitoring are limited, and citizen science can complement data from local management agencies. Here, citizen science efforts document the ecological status of a subtropical reef, Flinders Reef, Moreton Bay Marine Park, Australia. Ecological surveys, following Reef Check Australia and CoralWatch protocols, were conducted by 44 trained volunteers. Ten sites at Flinders Reef were surveyed at 5-10 m depth in autumn and spring. Additionally, underwater photos and depth surveys were integrated with satellite imagery to create a detailed habitat map. Coral cover across sites ranged between 14% and 67%. Branching corals dominated the site with 67% cover and showed 89% dissimilarity in coral community composition compared to other sites. Coral community composition was mostly explained by spatial variation, of which 16% was influenced by wave exposure. Observed reef impacts including physical damage, unknown scars and coral disease were three times lower than studies on more accessible reefs in Moreton Bay Marine Park. Invertebrate abundance was relatively low (6.65 individuals per 100 m^2^), with the most abundant groups observed being sea urchins (*Diadema* spp.), gastropods (*Drupella* spp.) and anemones. Butterflyfish were recorded at every site and were the most abundant fish group surveyed. Findings highlight the healthy condition of Flinders Reef, likely influenced by its offshore location and protection status as a ‘no-take’ zone. This study demonstrates that increasing the current 500m radius protection zone by a further 500m could double the protected area of coral, offering potential further conservation benefits. The findings resulting from the ecological data analysis and created benthic habitat map, provide an example of how citizen science based projects can assist marine park authorities and the public in ongoing conservation efforts.

## Introduction

Subtropical, high-latitude reefs occur in a transition zone between tropical and temperate regions. The mixing of warm and cold waters creates a unique climate in which marine communities comprise tropical, subtropical and temperate species [1-3]. Though subtropical coral community diversity is generally lower than on tropical reefs, coral cover can be comparably high in some locations [4].

High-latitude coral communities of eastern Australia, such as those in Moreton Bay, Queensland, are commonly dominated by generalist, stress-tolerant species that are seemingly well adapted to marginal environmental conditions [5]. Reef building coral taxa typical of tropical reefs are often absent, macroalgal cover often high, and communities are commonly structured by wave energy and exposure [6-9]. Like their tropical counterparts, these subtropical reefs are subject to climate change [5,10], as well as more localised anthropogenic stressors including pollution, eutrophication, overfishing, and other physical damage [3,11]. As sea surface temperatures increase, subtropical reefs are commonly promoted as potential refuges for the conservation of tropical reef species moving poleward under future climate change scenarios [10,12-14]. Despite the ecological value of subtropical reefs now and into the future, they typically receive less attention and are understudied compared to tropical reefs, which are highly recognised for their high biodiversity and importance to tourism [12,15].

The pressures of rapid population growth in the southeast Queensland region of Australia have been specifically highlighted for Moreton Bay [11], located in proximity to the greater Brisbane area of 2.3 million people [16]. To understand the potential impacts of ever-increasing pressures on coral communities in this region and deliver management strategies to ensure their longevity, the collection of long-term ecological monitoring data is essential [17]. Unfortunately, many subtropical reefs have limited long-term monitoring programs in place [15].

Citizen science programs that engage the community in data collection, analyses and reporting can provide scientific data to monitor changes and contribute to the development of effective environmental management strategies [15,18-20]. Global citizen science organisations such as Reef Check (http://www.reefcheck.org) and CoralWatch (https://www.coralwatch.org) empower citizens to carry out visual surveys of coral health, benthic habitat composition, anthropogenic impacts and invertebrate and fish biodiversity on coral reefs. This information is collated into global publicly-accessible databases that contribute to research, management and conservation practices [21-24]. In addition to generating scientific data, citizen science programs improve community knowledge about ecosystem function and threats, thus enhancing public stewardship of those ecosystems [18,22,25].

Flinders Reef is a relatively small subtropical reef located at the northern entrance to Moreton Bay. The reef is protected as a Marine National Park Zone (also referred to as a green zone, i.e., a ‘no-take’ area where extractive activities like fishing or collecting are not allowed without a permit) within the Moreton Bay Marine Park. Moreton Bay provides habitat for many marine species including over 1,600 invertebrates, 125 coral species, 9 species of dolphin, migrating humpback whales, manta rays, grey nurse sharks, leopard sharks, and large herds of dugong [26]. Reef Health Impact Surveys [23] are carried out at Finders Reef by Queensland Parks and Wildlife Services intermittently, and CoralWatch surveys [22,27] are conducted opportunistically by visiting scuba divers. Since 2007, Reef Check Australia (www.reefcheckaustralia.org) has coordinated trained volunteers to carry out annual ecological surveys at Flinders Reef, limited to four sites due to access restrictions [28]. Research conducted by academic professionals at Flinders Reef has primarily focused on specific taxonomic groups such as fish [29], corals [4,30,31], sponges [32,33] and molluscs [34].

The objectives of this study were to undertake a comprehensive ecological assessment that included ecological surveys and baseline habitat mapping through citizen science based surveys of subtropical Flinders Reef, and to compare the spatial distribution of coral community composition with wave exposure. This study, known as the Flinders Reef Ecological Assessment (FREA), provides a detailed characterisation of the community composition at Flinders Reef in terms of benthic coverage, reef impacts, abundance of fish and invertebrate indicator species, and coral health status for 10 sites, accompanied by the first detailed habitat map of the reef. Habitat maps provide an important management tool, but currently the existing map of Flinders Reef is limited to an outline of the exposed sandstone platform without any spatial description of benthic composition.

Ecological survey protocols were based on globally recognised Reef Check and CoralWatch survey methods and were conducted by the citizen scientists of The University of Queensland Underwater Club (UniDive).

## Materials and Methods

### Study location and site selection

Flinders Reef is located on a small sandstone platform (6.5 ha) three nautical miles north of Moreton Island in the northern part of Moreton Bay Marine Park, southeast Queensland, Australia (26° 58.715’ S, 153° 29.150’ E) (Fig 1). A 500 m radius zone provides protection as a green zone under a no-take, no fishing, no collecting or anchoring policy since 2009 (Fig 1). The area beyond the green zone is designated as a Conservation Park Zone with eight moorings that are accessible to recreational boats for e.g., diving and fishing activities.

**Fig 1.**
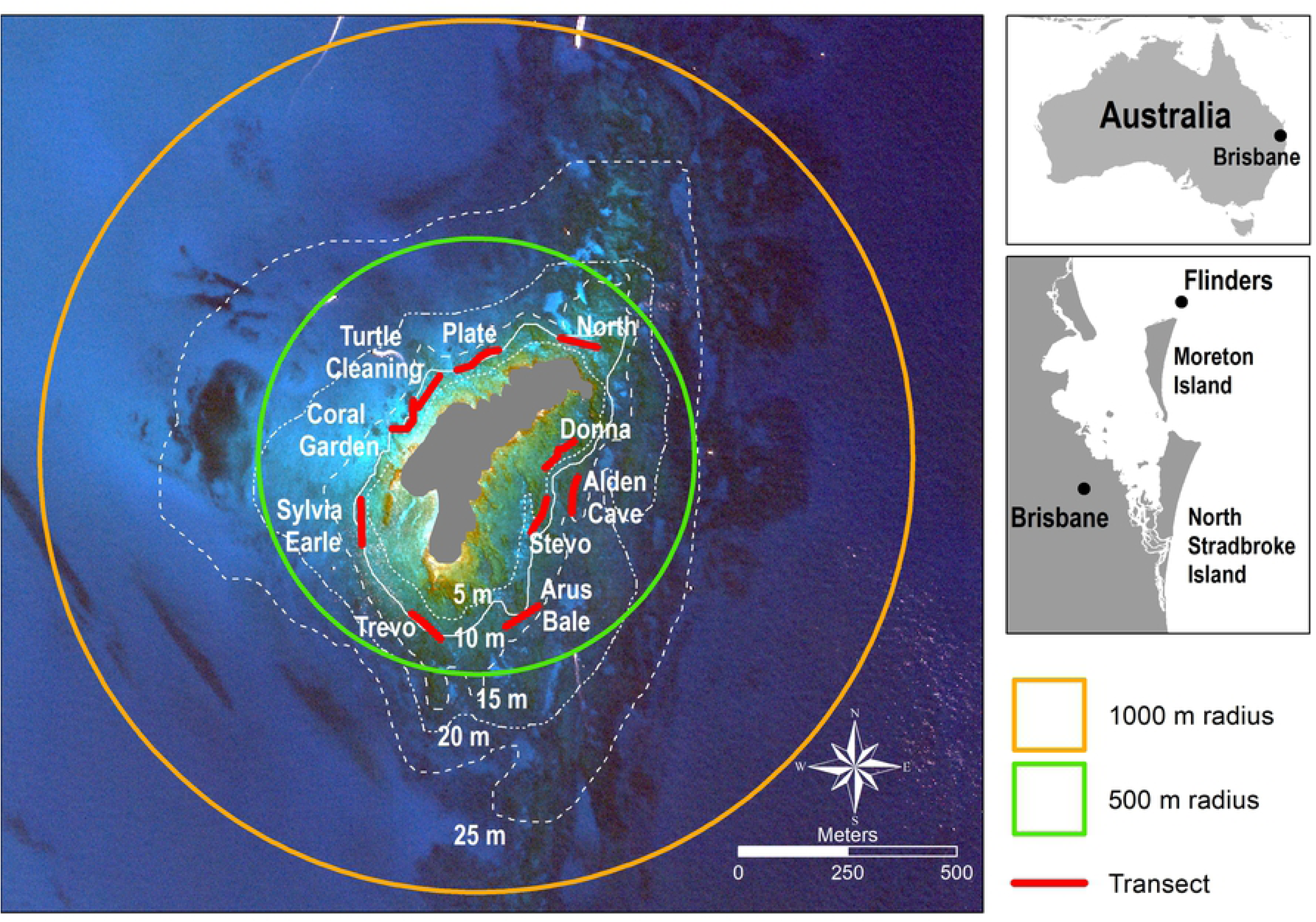
Satellite image of Flinders Reef with the approximate transect location and direction indicated in red lines (site names in white). The Marine National Park “Green” Zone (500 m radius) where no fishing or anchoring is allowed is designated by the green line. Orange line (1000 m radius) represents the suggested extension of the Green Zone (see discussion). The four Reef Check Australia long-term monitoring sites are Turtle Cleaning, Coral Garden, Plate and Alden Cave, respectively Turtle Cleaning Station, Coral Gardens, Alden’s Cave and Plateland in Reef Check Australia reporting. Source image: WorldView 2 image Digital Globe (2017), 2 m × 2 m pixels.

Due to the offshore location of Flinders Reef, access is limited and it is partly protected from nearshore environmental impacts such as poor water quality [3], but remains subject to potential climate change influences and pressures from direct use. Additionally, the offshore location likely promoted the development of its rich coral community, comprising 125 species [1,4,5,31]. Flinders Reef is on the southern distribution range of many tropical coral and fish species including *Acropora* spp. and Labridae [3,5,9].

Ten survey sites were established at 5-10 m depth within the green zone (Fig 1). Four of the ten sites have been visited annually since 2009 by Reef Check Australia: Alden’s Cave, Coral Gardens, Turtle Cleaning Station and Plateland. The ten sites were chosen as they represent a gradient in wind speed and direction, and wave height around the sandstone platform. Surveys were conducted in the Australian spring (March) and autumn (September) to capture any potential seasonal changes in marine communities. The site named Arus Bale was surveyed in autumn only due to weather conditions.

### Citizen science expertise and training

Approximately 100 UniDive members participated in the FREA citizen scientist project, mostly students, staff or alumni within the university. UniDive has a long history of award-winning citizen science projects in southeast Queensland [15,26,35,36]. Before any surveys were carried out, participants conducted academic and practical training in the ecological survey methods (Reef Check Australia and CoralWatch) and mapping survey methods provided by other members whom are experts in the field. To partake in field surveys, participants were required to be certified rescue (or equivalent) divers and qualify as Reef Check Australia divers by achieving a score of ≥85% on a theory exam, ≥95% on an in-water species identification exam and passing a practical in-water survey skills test. Eventually, 44 divers conducted the surveys and mapping, and ongoing training and quality control was conducted by the trainers throughout the project’s duration. Collected survey data underwent quality control for errors and inconsistencies via reviews of datasheets in the field and during data entry. If discrepancies were identified, recorded data was compared to survey photographs taken by the divers.

## Data collection

### Baseline benthic habitat mapping

The first detailed benthic habitat map of Flinders Reef was created following a previously-used protocol that involved delineating features visible in high spatial resolution satellite imagery based on colour and texture [15,26]. Habitat types were identified by overlaying the georeferenced field data onto satellite images. The georeferenced field data included: 1) water depth collected by boat echo sounder or diver; 2) feature surveys where divers would identify significant geological or ecological features present underwater; and 3) georeferenced photo quadrates representing a 1 m^2^ benthic footprint, captured at 1 to 2 m intervals along the seabed while the photographer towed a surface GPS recording the position [37].

### Baseline ecological and reef impact surveys (Reef Check Australia)

Ecological and reef impact surveys [21,38] consisted of visual surveys of substrate, reef health impact, invertebrate and fish indicator categories (S1 Table). At each site, surveys were conducted along one transect consisting of four 20 m segments, each separated by a 5 m gap. Substrate surveys included a point intercept sampling method where the substrate category was recorded at 0.5 m intervals along each transect, resulting in a percentage cover per site. Visual census of invertebrate and signs of reef impact categories were recorded along a 5 m belt by following a ‘U-shaped’ pattern along each 20 m segment, covering a total of 100 m^2^ per segment. The visual fish census survey divers recorded any fish categories observed within an imagined 5 × 5 m tunnel along each segment, i.e., 100 m^2^ area per segment. To ensure standardisation of the method and a constant detection probability, reef health impact, invertebrate and fish surveyors spent 7-10 minutes in each segment. Recognising the subtropical nature of Flinders Reef, existing methods were slightly modified by adding additional indicator species. For the substrate category, corallimorphs were added and for the fish surveys, the following groups were added: blue groper (*Achoerodus viridis)*, spangled emperor *(Lethrinus nebulosus)*, other emperors (*Lethrinidae*) and morwongs *(Cheilodactylus fuscus* and *C. vestitus*). For further analysis, indicator categories were consolidated into larger groups for visualisation purposes (S1 Table).

### Coral health surveys (CoralWatch)

Coral health data was collected using CoralWatch protocols [22,27], which involve assessing the colour of coral as an indicator of coral health by comparing the colour of a live coral with a pre-calibrated colour chart. The colours in the chart represent the health of the coral, i.e., the darker the coral colour, the healthier the coral is. The surveyor swam along the 5 m wide belt transect and assessed the coral colour and growth form of 5 randomly selected coral colonies per segment, totalling 20 coral colonies per site.

### Wave exposure

Wave height at Flinders Reef was determined using a third-generation wave nearshore model (Simulating WAves Nearshore, SWAN) [39]. Wave inputs for the SWAN model were based on the 1976-2017 wave record from the Brisbane wave rider buoy operated by the Queensland Department of Environment and Science. The wave rider buoy is located in deep water, east of North Stradbroke Island and southeast of the field site. The wave conditions during the 41-year period had a significant wave height *(Hs)* of 1.67 m, a wave period (*T*) of 9.43 s, and a wave direction (Dir) of 120.7°. Bathymetry for the SWAN model was generated from the Australian bathymetry and topography 2009 data set produced by Geoscience Australia [40]. A near neighbour interpolation method was used to convert the 9 arc second Ausbathy grid to a 50 × 50 m bathymetric grid for Flinders Reef and surrounding region, including the north and east coast of Moreton Island. The default parameters in SWAN were selected for wave modelling. Values of significant wave heights for each site were extracted based on the centre coordinates of each transect in a Universal Transverse Mercator (UTM) coordinate system.

## Data and statistical analyses

### Data manipulation

Differences between autumn and spring surveys were tested using a Student’s t-test based on the overall mean of measurements for the four survey types, i.e., substrate, reef impact, invertebrate and fish. No significant differences across seasons were found, therefore, measurements were averaged across seasons (S2 Table). Measurements per survey type were then averaged across seasons and the four segments representing the site. The majority of surveyed reef impacts specifically affect corals and hence an area with high coral cover is likely to have inherently greater impact abundance. To allow comparison of reef impact abundance between sites of varying coral cover, reef impact abundance was normalised to percent hard coral cover. Impact, invertebrate and fish group abundance was calculated per 100 m^2^.

### Multivariate analyses to assess coral community composition

Further analyses focusing on coral community composition were based on only the seven hard coral and four soft coral categories monitored (S1 Table), as responses in coral community composition are commonly driven by environmental variables such as wave exposure [6,7]. A hierarchical clustering approach was used to determine dissimilarities in coral community composition between sites. Distances were measured using a Bray-Curtis dissimilarity matrix and a complete linkage cluster aggregation method was performed in order to form clusters. The significance of the clusters was tested using a cluster-based permutation test based on 999 permutations and an error-level of 0.05. Non-metric multidimensional scaling (nMDS) ordination was then used to locate dissimilarities in coral community composition across sites based on the four segments surveyed. The nMDS is displayed in a two-dimensional space and shows Bray-Curtis dissimilarity distances between sites. A third multivariate analysis was performed in order to quantify the influences of sites and wave exposure on coral community composition at Flinders Reef. A permutational multivariate analysis of variance (PERMANOVA) was used based on Bray-Curtis dissimilarity distances in coral community composition with segments nested within sites and wave exposure formulated as fixed effects. Significance was tested based on 999 permutations at a 0.05 error-level. Multivariate statistical analyses were performed using the R packages “clustsig” [41] and “vegan” [42] within R version 3.2.2 software [43].

Values of wave exposure at sites were correlated with coral communities across Flinders Reef. The Pearson product moment correlation coefficient and associated p-value indicated the direction of the relationship between wave exposure and proportion of coral categories at a site and significance at 0.05 error-level. Two levels of wave exposure were calculated using the median values of wave height across all the sites. These levels were then combined with the outputs of the nMDS to examine the influence of wave exposure on the observed clustering of coral communities.

### Coral health chart analysis (CoralWatch)

At Flinders Reef, the lightest and darkest colour score was measured for a total of 378 coral colonies. The average colour score at Flinders Reef and per site (± standard error (SE)) was calculated by pooling the two seasons (autumn and spring).

## Results

### Baseline benthic habitat mapping

The georeferenced habitat map created for Flinders Reef describes substrate type, water depth and significant features (Fig 2) [44]. Prominent mapped features at sites are vast branching hard coral beds at Coral Garden and large plate corals with diameters up to ∼2 m at 10-15 m depth near Plate, and in deeper water south of Alden Cave and Trevo. Encrusting and plate corals were observed mostly on the southeastern side, with branching hard corals and soft corals on the western side. *Asparagopsis* sp. was the dominant macroalgae observed at Flinders Reef, however, macroalgae *Laurencia* sp. was more abundant in deeper waters (>15 m). Rock and rubble surfaces not covered by coral were covered by macroalgae or turf algae. Sandy areas were predominantly found in deeper waters (>15 m).

**Fig 2.**
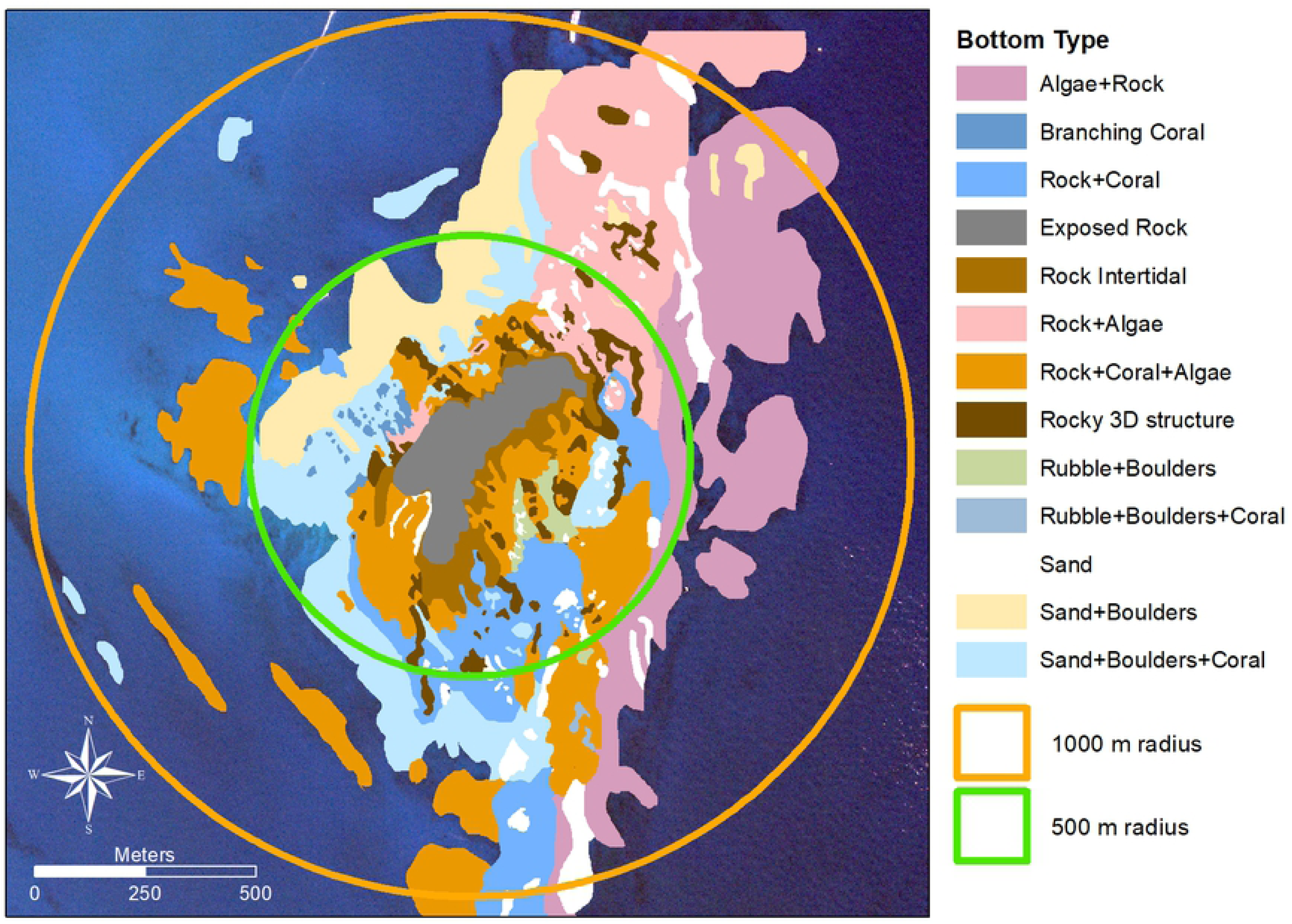
Detailed habitat map of prominent substrate types for Flinders Reef, southeast Queensland, Australia. Marine National Park “Green” Zone (500 m radius, green line) where neither fishing nor anchoring is allowed could be extended with an additional 500 m buffer zone (1000 m radius, orange line) where no anchoring would be allowed. This would result in a two-fold increase in protected area for bottom types that contain coral communities; and a three-fold increase of areas that include hard rock substrate.

### Ecological baseline

There was no significant difference in the overall mean of measurements for the four survey types between autumn and spring at Flinders Reef (p > 0.050, S2 Table). Therefore, results were pooled across the two seasons.

Across all survey sites, the most common group identified in the substrate composition survey was rock with an average cover of 37.0% (per 100 m^2^), followed by hard coral (33.3%) and soft coral (10.0%). The maximum and minimum hard coral cover was recorded at Coral Garden (66.9%) and Plate (14.0%), respectively (Fig 3A).

**Fig 3.**
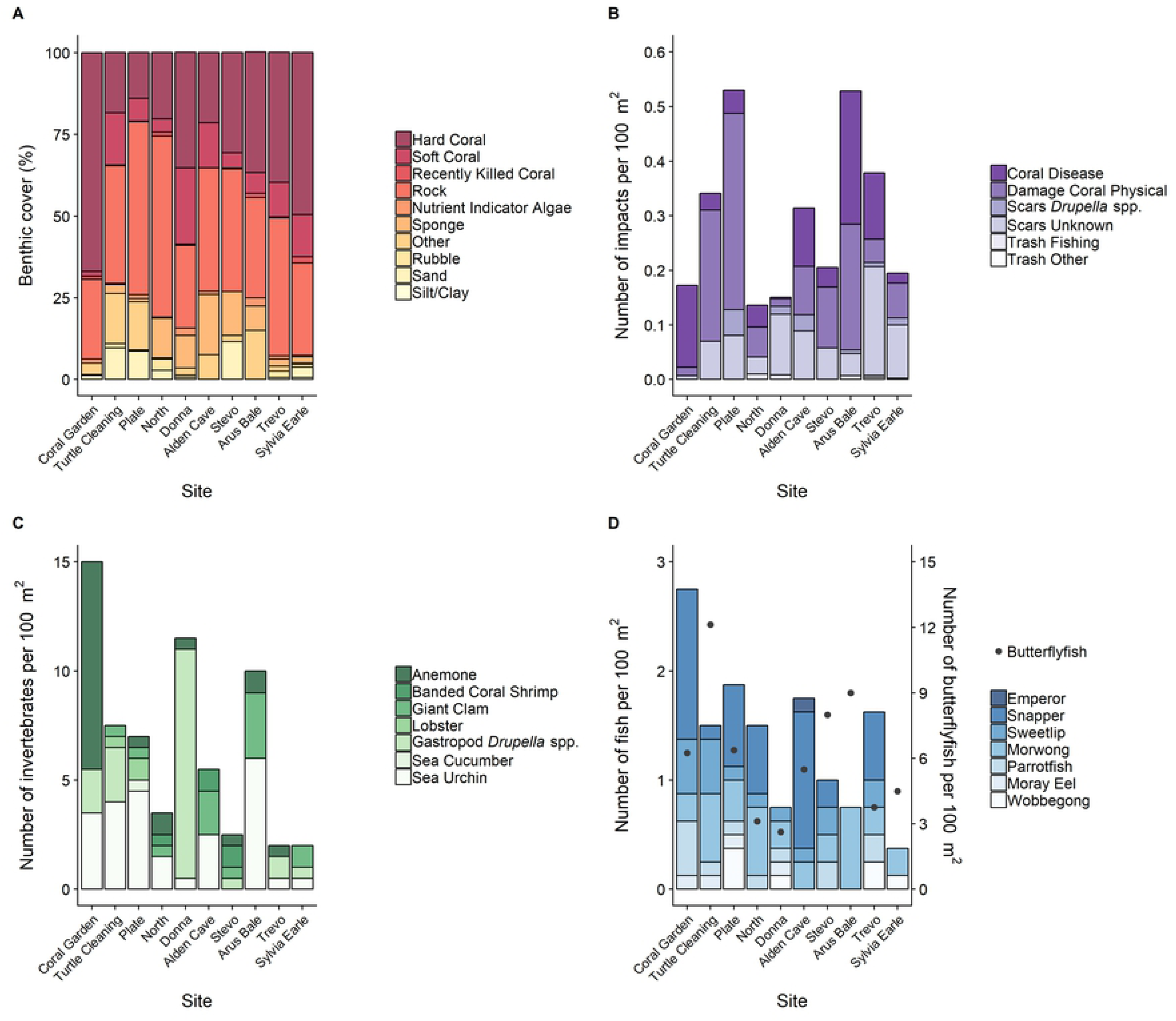
Overview of the major ecological groups recorded during the surveys per site. (A) Benthic groups expressed as percentage cover per site. (B) Average number of impacts per site per 100 m^2^ normalized for hard coral cover. (C) Average number of invertebrates found per site per 100 m^2^ and (D) average number of fish per site per 100 m^2^ with the average number of butterflyfish displayed on the secondary y-axis (dots). Groups were based on surveyed ecological categories and absent categories/groups were omitted from the panels.

Overall, the number of reef impacts detected was low (Fig 3B). The most common impacts observed were coral physical damage (average of 0.12 occurrences per 100 m^2^) followed by coral disease and unknown scars (both averaged 0.08 occurrences per 100 m^2^). Turtle Cleaning and Arus Bale exhibited the greatest prevalence of impacts, driven by coral physical damage and also coral disease at Arus Bale. Three reef impact categories were not observed: crown-of-thorn starfish (*Acanthaster planci*) scars, coral damage due to boat anchor or due to dynamite. Coral health chart colour indicator surveys demonstrated a healthy reef with an average colour score measured at Flinders Reef of 3.9 ± 0.07 standard error (SE). The highest average colour score was measured at Trevo (mean ± SE, 4.4 ± 1.8) and lowest score at Arus Bale (2.7 ± 0.22).

The overall number of invertebrates seen was relatively low (average of 6.65 individuals per 100 m^2^, Fig 3C). Nevertheless, the presence and abundance of indicator invertebrate categories varied between survey sites with the most diverse site being Plate, i.e., 5 out of 14 recorded taxa observed (Fig 3C). The most abundant invertebrate groups were sea urchins (especially *Diadema* spp.), gastropods (*Drupella* spp.) and anemones with on average 2.35, 1.70 and 1.35 individuals per 100 m^2^, respectively. Coral Garden exhibited the highest number of invertebrates, primarily anemones (9.50 per 100 m^2^), and the highest abundance of *Drupella* spp. was found at Donna (10.50 per 100 m^2^). Crown-of-thorn starfish (*Acanthaster planci*), gastropods triton (*Charonia tritonis*) and trochus (*Tectus niloticus*), sea cucumbers prickly greenfish (*Stichopus chloronotus*) and prickly redfish (*Thelenota ananas*), and pencil sea urchins (*Heterocentrotus mammillatus* and *Phyllacanthus parvispinus*) were included in the surveys, but not observed at any of the sites.

Fish community composition was largely dominated by butterflyfish which were recorded at each of the ten sites (Fig 3D), and a total of 524 individuals were counted during the surveys. On average, 6.12 butterflyfish were recorded per 100 m^2^ ranging from 2.62 at Donna to 12.10 at Turtle Cleaning. The second-most dominant fish group was snapper with 40 individuals recorded at seven sites and an average of 0.50 fish per 100 m^2^, followed by morwong (0.39), sweetlip (0.20) and parrotfish (0.15). A substantial number of fish categories were not recorded during surveys, i.e., coral trout, Queensland grouper (*Epinephelus lanceolatus*), barramundi cod (*Cromileptes altivelis*), other groupers, blue groper (*Achoerodus viridis*), humphead wrasse (*Cheilinus undulates*), bumphead parrotfish (*Bolbometopon muricatum*) and pink snapper (*Pagrus auratus*).

### Coral community analysis

There was 89% dissimilarity found in coral community composition between Coral Garden (Cluster 1) and the remaining sites (Cluster 2) (p = 0.016, Fig 4A). Within Cluster 2, a dissimilarity of 58% separated the northwestern sites (Turtle Cleaning and Plate) from the others; however, this clustering pattern was not significant. The nMDS ordination plot (stress = 1.74%) indicated that Cluster 1 was dominated by branching corals (Fig 4B) representing 64.1% of the substrate cover at Coral Garden according to the ecological surveys. Cluster 2 comprised a mix of coral indicator groups. Coral community composition at the northwestern sites, Turtle Cleaning and Plate, was characterised by plating and foliose hard corals. In comparison, sites on the eastern side of Flinders Reef, Alden Cave, North and Trevo, were characterised by encrusting hard coral (Fig 4B).

**Fig 4.**
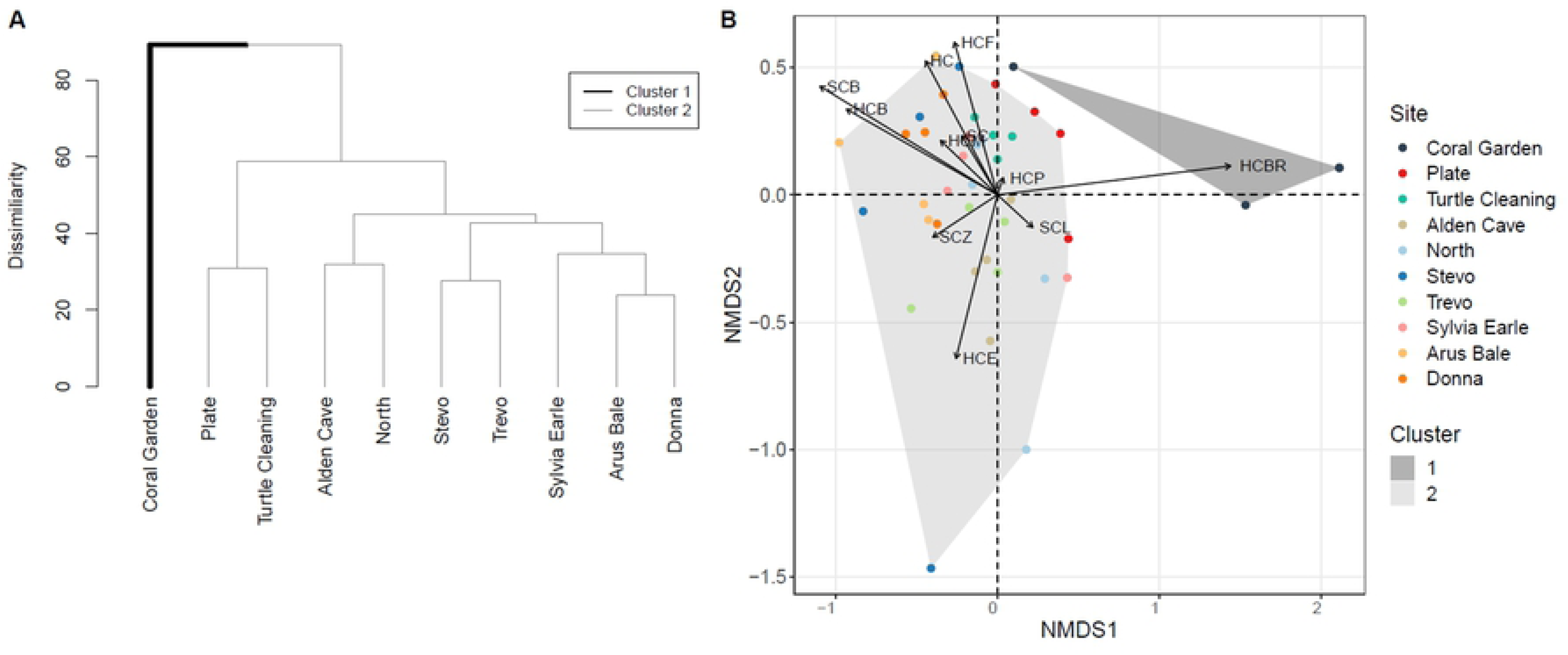
Clustering of sites according to coral community composition. (A) Dendrogram depicting the significant hierarchical clustering of coral community composition of the surveyed sites around Flinders Reef and associated dissimilarity values. (B) Non-metric multidimensional scaling (nMDS) plot using Bray-Curtis dissimilarities (stress = 1.74%), illustrating dissimilarities between the four segments per site, averaged across seasons. Arrows indicate the coral community groups driving these dissimilarities. Cluster 1 and 2 identified in the dendrogram (Panel A) are represented by grey polygons in Panel B.

### Wave exposure and community composition

Sites located on the northwestern side of Flinders Reef were the least exposed to waves. Significant wave height was 0.9 m for Turtle Cleaning and Plate, and 1.2 m at Coral Garden (Fig 5A). Wave heights for the seven remaining sites varied between 1.5 and 1.6 m (Fig 5A). The median significant wave height across all sites was 1.54 m, separating the less exposed sites (located west to north of Flinders Reef) from the more exposed sites (east to south), forming two groups. There was a positive relationship between wave exposure and the proportion of encrusting corals (HCE, p < 0.001) and soft coral zoanthids (SCZ, p = 0.008), and a negative relationship with leathery soft coral (SCL, p = 0.021) (Fig 5B). Fragile hard corals (HCF, HCP and HCBR) and soft corals (SC) were associated with lower wave exposure, while more robust hard coral types (HCM, HC and HCE) were associated with higher wave exposure (Fig 5C).

**Fig 5.**
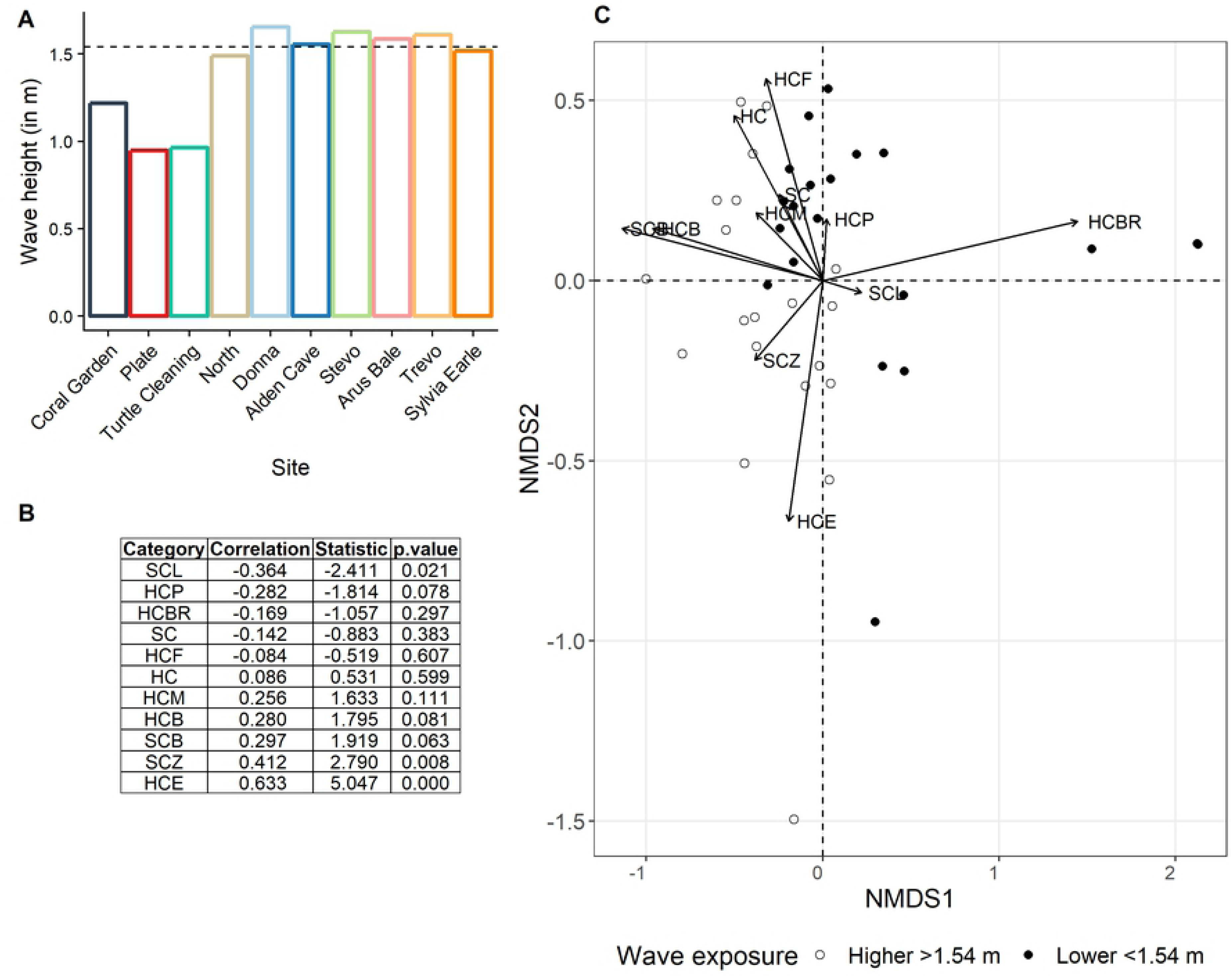
Wave height and observed relationship with coral community categories at each site. (A) Values of wave height extracted from the SWAN model at each site. The dotted line shows the median value of 1.54 m, used to separate sites with lower and higher wave exposure into two groups. (B) Correlation values and associated significance between distinct coral categories and wave height at Flinders Reef. (C) Combines wave exposure for the two groups with the non-metric multidimensional scaling plot (Fig 3C). Each dot represents a 20 m survey segment within a site.

There was a significant contribution of site (p < 0.001) and wave height (p < 0.001) to coral community composition (Table 1). A total of 47.0% and 15.6% of the variability was explained by site and wave exposure, respectively.

**Table 1.** Contribution of site and wave height to coral community composition.

| Source | df | SS | MS | F | R<sup>2</sup> | P-value |
| --- | --- | --- | --- | --- | --- | --- |
| Wave height | 1 | 0.9774 | 0.97743 | 12.5046 | 0.15594 | <0.001 |
| Site | 8 | 2.9458 | 0.36822 | 4.7108 | 0.46996 | <0.001 |
| Residuals | 30 | 2.345 | 0.07817 |  | 0.37411 |  |
| Total | 39 | 6.2682 |  |  | 1 |  |

Summary PERMANOVA output including coral community composition as response variable, and wave height and site as fixed effect explanatory variables.

## Discussion

FREA was facilitated by more than 100 volunteer citizen scientists who recorded and revised the ecological and geographical characterisation of Flinders Reef in Moreton Bay Marine Park. Together, the results of the ecological and mapping surveys report on: the distribution of habitats; coral community composition; ecosystem condition; and biodiversity via indicator species, to produce a comprehensive ecological assessment of this subtropical reef. This study highlights the remarkable coral cover at this subtropical reef, with some sites having comparable cover to the Great Barrier Reef [45], which is consistent with a previous subtropical reef study [9]. There was also a clear zonation of coral community composition, notably influenced by site location and wave exposure. Coral Gardens is a unique site at Flinders Reef with high branching coral cover. Soft corals and fragile hard coral morphologies such as extensive branching coral beds were associated with the northwestern reef sites, characterised by lower wave exposure, while more robust hard coral morphologies were found on the exposed eastern and southeastern sites. Branching hard corals are more susceptible to damage from waves and storm events [46], which may explain the dominance of fragile branching coral on the sheltered side of Flinders Reef. The observed zonation patterns align with previous documentation of Flinders Reef [4], as well as other studies on the influence of wave exposure on high-latitude coral reef community assemblages [6,47]. In addition to wave exposure, coral community composition may be influenced by the intensity and regularity of disturbance [48], the depth at which the coral community is located [49], and patterns of recruit settlement [50].

While physical damage, unknown scars and coral disease were recorded at all monitoring sites, overall impacts at Flinders Reef were three times lower than those observed for more accessible reef locations in Moreton Bay at Point Lookout [15]. Coral health chart surveys also indicated healthy corals with no signs of coral bleaching. Earlier studies, before the establishment of the green zone, reported anchor damage at Flinders Reef [4]. The lack of anchor damage in the present study suggests that the installation of moorings and establishment of a green zone (with no anchoring) is effective in protecting the reef from damage. The 500 m radius green zone around Flinders Reef is a ‘no-take’ area and the site is relatively far from the mainland (1-2 hours travel by boat) compared to other sites within the Moreton Bay Marine Park, all of which may limit visitation and use. Yet, higher levels of coral damage were recorded at the most popular dive locations around Flinders Reef, which have the highest cover of branching coral. This could reflect damage from scuba dive tourism or other site visitation and/or the fragility of branching coral compared to other coral morphologies [46]. The effectiveness of the green zone is also supported by the lack of fishing lines found during impact surveys of the present study. However, there are anecdotal reports of fishing within the protected area, and close surveillance of poaching activities can be made difficult by the remoteness of the location.

A relatively low abundance of target invertebrates was found across sites, which is generally consistent with long-term Reef Check Australia findings [51,52]. While there were many closely-related and functionally-equivalent invertebrates present on our transects, these were non-target species according to the survey protocol [38,53]. The distribution of invertebrates varied spatially. The abundance of gastropods (*Drupella* spp.) did not seem to correlate with the cover of hard coral nor with recorded abundance of *Drupella* scars. Corallivorous gastropods may have formed isolated aggregations within the surveys while the overall distribution of gastropods (*Drupella* spp.) might be low, as observed in other coastal waters [54]. The high abundance of anemones at Coral Garden might be related to wave exposure. A previous survey of subtropical anemones found that abundance was significantly higher on leeward reef sites compared to those that were more exposed [55]. Though pencil sea urchins (*Heterocentrotus mammillatus* and *Phyllacanthus parvispinus*) are known to be present at Flinders Reef [56-59], they were not recorded in this study. This may be attributed to low abundance, seasonal differences and/or local movement of the animals.

Butterflyfishes dominated the fish community and were observed at each survey site, in highest abundance at Turtle Cleaning. A consistently high abundance in butterflyfishes has been observed in other similar locations to Flinders Reef [51,52]. Many butterflyfishes are corallivorous fish and mainly target hard coral with some species preferring soft coral polyps as a food source [60]. They have distinct prey preferences that can be specific to one coral species, genera or growth morphology [60], which can limit their abundance and distribution. However, without knowledge of the butterflyfish species at each site, we cannot conclude that differences in butterflyfish abundance across sites is related to the hard or soft coral cover or diversity. Fish community composition and abundance is often influenced by live coral cover and structural complexity [61,62], which may explain the highest fish abundance observed at Coral Garden. However, aside from butterflyfishes, fish abundances were relatively low. Most target fish groups have larger home ranges and may prefer deeper areas away from the currents, surge and exposure of the rock platform. Additionally, parrotfish and grouper abundance may have been underestimated due to the inclusion of only larger sized individuals (surveys only included parrotfish >20 cm and grouper >30 cm). Smaller juvenile fish are known to use shallower reef areas as nurseries and have smaller home ranges compared to adult fish [63,64].

This study highlights the important role of citizen science in providing information to enhance current monitoring outcomes through improved site selection and expanded indicator categories. The habitat mapping and ecological benthic data results suggest that there may be benefits of regularly monitoring additional survey sites, e.g., Sylvia Earle, to produce a fully representative sample of the habitat diversity at Flinders Reef. The habitat mapping approach is accessible to almost any citizen science project and provides high quality data. Although it requires thorough training, as well as the open source GIS, and the acquisition of specific software, satellite imagery and standard underwater camera equipment, we hope this method will become more widely applied in coral reef surveys by providing detailed methodology protocols [this study, 26]. In regards to indicator species, in Reef Check Australia protocols these have been selected for broad geographic coverage with a focus on tropical species [53]. Including broader functional survey categories for fish and invertebrates, as well as including all fish sizes in future surveys may improve ecosystem health monitoring on subtropical reefs. Additionally, inclusion of tropical species in these surveys will be increasingly important to detect ‘tropicalisation’ of subtropical marine environments, i.e., the movement of tropical species poleward [5,10,65,66]. The value of these species as indicators will strongly depend on their dispersal potential.

The current green zone comprises a 500 m radius circle from the centre of the Flinders Reef sandstone platform. The ecological assessment and habitat mapping provide a detailed description of the benthic composition of Flinders Reef and highlight deeper reef habitats that are currently excluded from the green zone. This may prompt consideration for expansion of the green zone to a circular area of 1000 m radius. Such an expansion would result in: inclusion within the green zone of all areas mapped with coral communities to a depth of 25 m; a two-fold increase in surface area for bottom types that contain coral communities; and a three-fold increase of areas that include hard rock substrate (Fig 1 and 2, orange polygon), which is required for coral settlement and post-settlement survival [67]. Expansion of the green zone may increase opportunities for corals to settle and subsequently enhance coral growth and abundance of reef-associated fish. Green zones have been shown to enhance recreational fishing opportunities outside of the protected area through increased fish biomass and abundance [68], benefiting both fishermen and the ecosystem.

## Conclusion

The FREA citizen science project brought together a range of community members and partners to document the healthy status of a high-latitude reef, Flinders Reef, with coral cover at some sites comparable to locations on the Great Barrier Reef. This study highlights the value of citizen science as an approach to complement traditional scientific and management approaches, as well as engaging local community members to learn about and take active steps to care for local environments. The volunteer support of over 10,000 hours, 500 dives, 44 trained divers and community-funded investment falls outside the standard resource capability of marine park management. The project also facilitated a number of opportunities for local divers to learn more about reef ecology, and enhanced community support, understanding and project ownership through a range of communication tools including a technical report, coffee table photo book, posters, television segments and community events. The study offered a platform for constructive discussions and applications around the monitoring, management and stewardship of Flinders Reef into the future.

## Acknowledgements

We would like to acknowledge the 44 core survey divers and many additional UniDive volunteers who spent >10,000 hours on the FREA project. Financial support was given by the Queensland Parks and Wildlife Services, Honourable Dr Steven Miles – former Minister for Environment and the Great Barrier Reef, Solar School, Healthy Land and Water, and those who supported the ING Dreamstarter crowdfunder. In-kind support provided by Point Lookout Scuba Dive Charters (Ken Holzheimer); The University of Queensland Boating and Diving; Moreton Island Adventures; Tangatours; Queensland Parks and Wildlife; Geoimage; Dr Ian Tibbets, The University of Queensland; Aquatic Centre, The University of Queensland; Reef Check Australia and CoralWatch.

## Supporting information

**S1 Table: Reef Check Australia (RCA) indicator categories per survey type (i.e., substrate, impact, invertebrate and fish) consolidated to Flinders Reef Ecological Assessment (FREA) groups for data collection and statistical analysis.** Substrate and fish survey RCA categories were modified for the FREA citizen science project.

**S2 Table: Results seasonal survey comparison.** Summary Student’s t-test output based on the overall mean of measurements for the four survey types, i.e., substrate, impact, invertebrate and fish.

